# Unraveling novel insights into dual-species cariogenic biofilm formation: a comparative analysis on natural vs artificial bioengineered dentin models

**DOI:** 10.1101/2025.05.24.655910

**Authors:** Javiera Ortiz, Simón Álvarez, Sebastian Aguayo

**Author notes:** Corresponding author: Dr. Sebastian Aguayo, PhD, School of Dentistry and Institute for Biological and Medical Engineering Pontificia Universidad Católica de Chile, E.

## Abstract

Dental caries is the most prevalent biofilm-associated disease affecting billions of people worldwide, including elderly individuals. Conventional biofilm study methods rely on human or animal-derived samples, posing challenges regarding accessibility, cost, and ethical considerations. While *in-vitro* systems offer a promising alternative, they often fail to replicate the structural characteristics of dentin, which play a crucial role in bacterial adhesion. To bridge this gap, a bioengineered dentin-like construct has recently been developed as a reproducible and accessible model for studying biofilm formation associated with dental aging. Therefore, this study aimed to assess dual-species *Streptococcus mutans* and *Candida albicans* biofilm formation on bioengineered dentin substrates and compare it to biofilm formation on natural human aged dentin. For this, *S. mutans* UA159 and *C. albicans* (ATCC 90028) were co-cultured on bioengineered and natural dentin slabs, and polymicrobial biofilm formation and EPS production were characterized via high-resolution confocal laser scanning microscopy. Following biofilm formation, image processing was conducted using COMSTAT software to determine biofilm growth parameters. Additionally, fluorescence intensity was quantified via microplate readings, and cell viability was assessed using a Live/Dead viability kit. Overall, results showed similar biofilm formation patterns between the bioengineered and aged dentin, with no significant differences found in biofilm physical properties or viability. These findings suggest that this bioengineered dentin system provides a reliable platform for studying biofilm formation in the context of dental aging, making it a valuable tool for investigating microbial adhesion and cariogenic biofilm development under controlled conditions, potentially facilitating future research in biofilm-related oral diseases.

**Importance:** Dental caries is one of the most common chronic diseases worldwide, which is driven by complex microbial biofilms formed on the tooth’s surface. However, existing models for studying these biofilms in the laboratory often rely on human or animal tissues, which are difficult to obtain and standardize, and present ethical challenges. In this study, we validate a bioengineered dentin-like model that accurately mimics the microarchitecture of aged human dentin, a key site for root caries in the elderly. By comparing biofilms formed by the clinically significant *Streptococcus mutans* and *Candida albicans* on both artificial and natural substrates, we show that the engineered model supports biofilm development under comparable parameters and enables detection of changes in microbial virulence. Overall, this platform provides a reproducible and scalable alternative for studying oral biofilms with potential applications in understanding disease pathogenesis, novel treatment testing, and integration into next-generation organ-on-a-chip systems.

## 1. Introduction

Caries is the most prevalent disease worldwide, affecting over 2 billion people (1). With a strong tendency toward aging populations (2,3), it remains necessary to continue promoting long-lasting oral health and developing specifically tailored preventive and therapeutic measures (2–4). In this context, understanding how host-pathogen interactions at the dental interface can lead to the development of dental caries is crucial to unveil new strategies. To date, the most employed methods for studying oral biofilm formation in the laboratory are *in-vitro* studies and animal models (5–7); however, these approaches can be costly, and the acquisition of animal or human samples may present significant bioethical and experimental challenges.

Currently, microfabricated and microfluidic devices enable dynamic *in-vitro* experiments with reduced costs and have been successfully applied to study the pathophysiology of various organs such as the heart, liver, and others (8–10). Building on this technology, “*tooth-on-a-chip*” models have been developed to investigate microbial, cellular, and biomaterial interactions by simulating different oral environments(11). One key aspect that can be studied using these models is biofilm adhesion(12); however, existing tooth-on-a-chip models still lack the incorporation of a critical factor for biofilm formation on dentin: substrate topography. Furthermore, while many static *in-vitro* models examine biofilm adhesion on hydroxyapatite disks that mimic the smooth enamel tissue surface, dentin tissue remains largely unexplored in such studies.

Dentin is the most abundant tooth tissue that has a tubular microarchitecture and is composed mostly of inorganic hydroxyapatite (HAP) but also around 20% of a type-I collagen organic matrix (13). Dentin exposure occurs due to enamel loss caused by caries, trauma, or gingival recession, which is estimated to affect two-thirds of the global population, with higher prevalence among older adults (14). Most importantly, dentin exposure allows colonization by resident oral biofilms and increases the risk of deep caries and root caries, which can lead to the loss of tooth vitality (15,16). Although the composition of dentin and its interaction with adhesive, restorative, or regenerative biomaterials are widely studied topics in dentistry, the influence of dentin architecture on microbial adhesion and subsequent biofilm formation is rarely considered in existing models (17).

Specific bacteria such as *Streptococcus mutans*, one of the most well-known cariogenic bacteria, is associated with the early formation of biofilms on dentin (18). It can adhere to collagen through specific collagen-binding adhesins such as Cnm, Cbm, WapA, and SpaP (19,20). Additionally, it has been observed to have a synergistic relationship with the opportunistic fungus *Candida albicans* (21,22), and together, they are linked to rapidly progressing caries in both infants and elderly root caries (23). Therefore, the *S. mutans-C. albicans* interaction may contribute to disease development, including in elderly patients. In this context, the accumulation of advanced glycation end products (AGEs) during aging has been associated with changes in collagen structure, as they act as cross-links between fibers, leading to alterations in the mechanical properties of collagen-rich tissues (24). In the specific context of dentin, this phenomenon results in increased stiffness of the collagen matrix (25). Furthermore, recent studies on bacterial adhesion to glycated dentin have shown enhanced biofilm adhesion and growth, which could partially explain the differences in dysbiosis observed in older adults and individuals with diabetes (26).

To address the lack of studies on bacterial adhesion to dentin, particularly in aged teeth, and to integrate this research into microfluidic devices, a bioengineered dentin-like construct mimicking the tubular architecture of dentin was recently developed by our group (27). These constructs can be experimentally glycated to simulate tooth aging, providing consistent and predictable substrates that ensure reproducible experimental results. Furthermore, its scaled-down composition reduces costs associated with reagent use and addresses the potential ethical concerns related to the acquisition and handling of human and animal materials.

Despite these experimental advances, the ability of these bioengineered constructs to allow the growth and development of oral biofilms, with a composition and structure comparable to those grown on natural older teeth, remains unclear. Therefore, the aim of this study was to compare early cariogenic biofilm formation on aged artificial dentin and aged natural human dentin in a sucrose-enriched environment, by employing a clinically relevant dual-species *S. mutans* and *C. albicans* biofilm.

## 2. Methods

### 2.1. Sample collection and experimental design

An *in-vitro* experimental study was conducted to evaluate biofilm formation on bioengineered and natural dentin substrates. Briefly, dual-species biofilms of *C. albicans* (ATCC 90028) and *S. mutans* (UA 159) were cultured at a 1:1 cell ratio for 24 hours in Brain Heart Infusion (BHI) supplemented with 1% sucrose, at 37°C and 5% CO₂. The natural dentin slabs were obtained from donated teeth (n=10), which were collected as waste material from patients (female or male) over 50 years of age without caries, following informed consent (ID: 21061500 Pontificia Universidad Católica de Chile). Briefly, caries-free teeth were selected and cleaned with 70° alcohol, then 300µm thick cross sections were obtained at the root level with a hard tissue microtome (Leica SP1600, USA). Approximately 20 slabs were collected per tooth. On the other hand, the bioengineered dentin-like substrates were constructed and glycated with methylglyoxal (MGO) as previously described(27). Biofilms were grown either with or without artificial saliva preconditioning, for which 100ul of artificial saliva (Sigma-Aldrich) was applied onto the substrates, incubated for 1 hour at room temperature, and removed prior to biofilm seeding. The experimental setup is illustrated in **Figure 1**.

**Figure 1:**
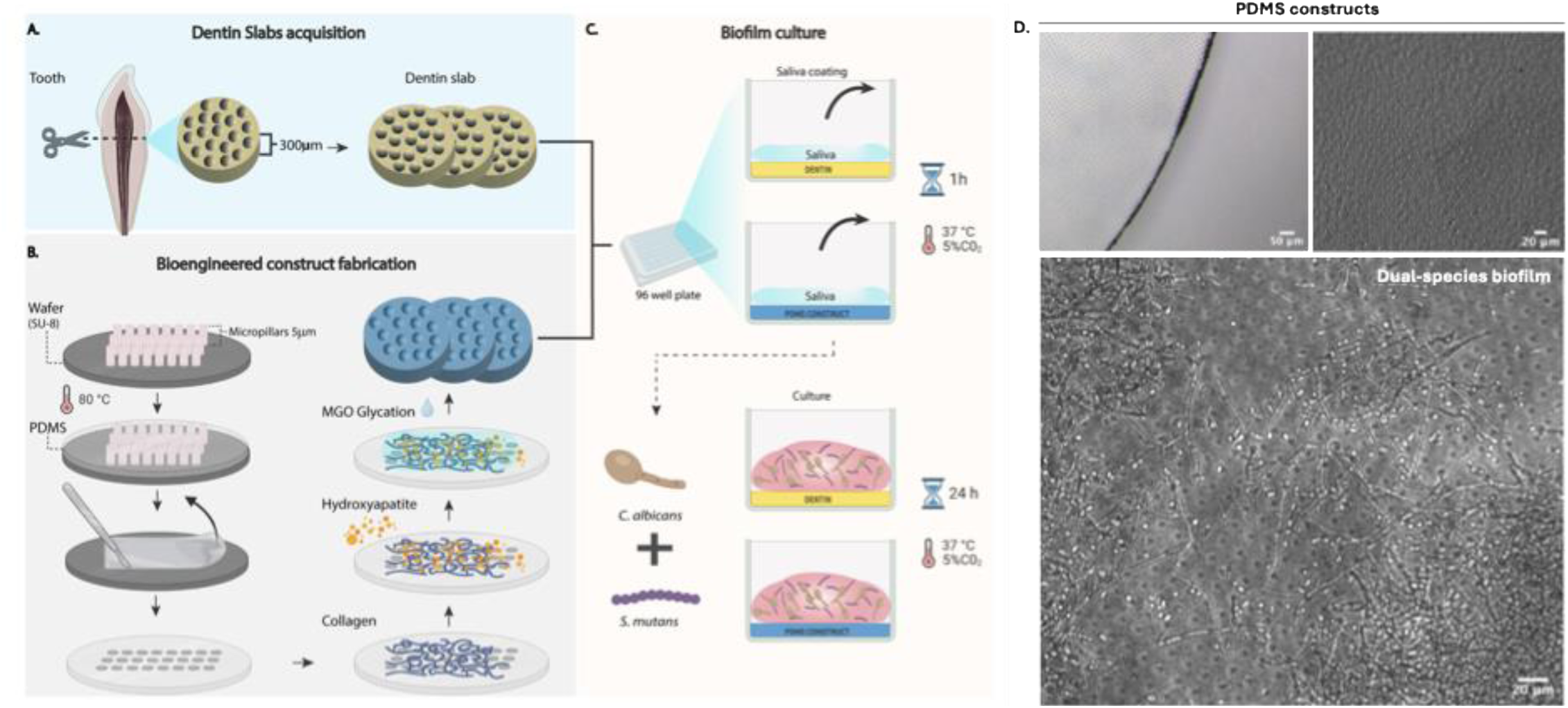
**Diagrammatic representation of the experimental setup regarding construction of the bioengineered dentin substrates and preparation of natural dentin specimen**. A. Tooth acquisition with cross-sectional cut; B. Bioengineered construct fabrication with PDMS and functionalization; C. Saliva coating and substrate culture with *S. mutans* and *C. albicans*. D. Optical microscopy images of constructed PDMS substrates (upper panels), and epifluorescence microscopy of a dual-species *C. albicans* and *S. mutans* biofilm grown on the biomaterial substrate (lower panel).

### 2.2. Biofilm viability assessment

To determine viability following surface growth, biofilms were stained using the LIVE/DEAD® BacLight™ Bacterial Viability Kit (Molecular Probes, USA), which employs a combination of 1.67 μM Syto® 9 and Propidium Iodide (PI) to differentiate live and dead/damaged bacteria, respectively. Stock solutions of PI (81845, Sigma; 20 mM) and SYTO 9 (S-34854, Invitrogen™, Thermo Fisher Scientific; 3.34 mM) were prepared and used according to the BacLight™ kit protocol. A 1:1 mixture of stains in PBS was prepared to achieve final concentrations of 30 μM PI and 5 μM SYTO 9. The stain mixture was added to cells harvested from surfaces by sonication (n=9). The samples were then incubated for 15 minutes in the dark at room temperature. As a control for dead staining, cells were treated with 70% ethanol for 1 hour prior to staining. After incubation, the supernatant was transferred to a 96-well microplate for spectrophotometric analysis (Synergy HT, Biotek).

### 2.3. High-resolution confocal fluorescence microscopy imaging of biofilms

The organization and structural architecture of the biofilms were assessed using simultaneous in situ labeling of bacterial cells and EPS with confocal laser scanning microscopy (CLSM). In summary, 1 µM Alexa Fluor 647-labeled dextran conjugate (Molecular Probes, USA; molecular weight: 10,000; absorbance/emission: 647/668 nm) was added to the culture medium during biofilm formation and development (1:1000). After 24 hours of incubation, excess stain was removed by washing once with PBS. The biofilms were then labeled with 2.5 µM SYTO 9 green-fluorescent nucleic acid stain in BHI (1:1000; Molecular Probes, absorbance/emission: 480/500 nm) for 30 minutes, followed by Calcofluor White (Sigma, M2R, 1 g/L, Evans blue, 0.5 g/L absorbance/emission:423-443 nm) in TSB (1:50) to stain fungal cells for 30 minutes. Careful PBS washes were performed after each staining step to remove excess reagent.

CLSM imaging was conducted using a Zeiss LMS 880 Axio Observer with an Airyscan detector. Images were acquired with a 40x water immersion objective lens (numerical aperture: 1.2), yielding an optical section thickness of approximately 1 µm. Excitation and emission wavelengths were set to 408–516 nm, 405–450 nm, and 633 nm for the respective fluorophores. The pixel size was set to 4084 x 4084, and image acquisition, processing and analysis was performed using ZEN Black 2.3 (ZEISS) software. A total of three samples were analyzed in triplicate, and the data were expressed as mean ± standard deviation. Statistical analyses were performed using GraphPad Prism Software 8.0 to compare differences between conditions.

### 2.4. Quantification of biofilm parameters by COMSTAT

Following structural characterization, CLSM image stacks were analyzed using the COMSTAT 2.1 image-processing software as previously described (28). Structural differences among the biofilms were evaluated by quantifying parameters such as roughness, total biomass, maximum thickness, and surface-to-biovolume ratio for total biofilm for each fluorescent channel. Furthermore, the three-dimensional architecture of the biofilms was visualized using the ZEN Blue Desk 2.3 software.

### 2.5. Quantification of Fluorophore Proportions in Dispersed Biofilms Using Microplate Fluorometry

To determine biofilm composition via fluorophore concentration after 24 hours of culture, stained biofilms as previously described were harvested using an ISOLAB ultrasonic water cleaner (maximum power: 60 W). Briefly, only substrate samples with adherent biofilm were transferred to sterile Eppendorf tubes containing 300 µL of PBS, vortexed for 30 seconds, sonicated for 1 minute, and vortexed again for 30 seconds (29). The supernatants were then transferred in triplicate to a 96-well microplate for fluorophore concentration measurement using a microplate reader (Synergy HT, Biotek). A total of three independent experiments were analyzed, and the data were expressed as mean ± standard deviation. Statistical analyses were performed using GraphPad Prism 8.0 to compare differences between conditions.

### 2.6. Quantitative PCR (qPCR) for virulence gene expression analysis

The expression of virulence-related genes in *S. mutans* and *C. albicans* was quantified using reverse transcription-quantitative polymerase chain reaction (RT-qPCR). In *S. mutans*, we analyzed *spaP* (encoding adhesin P1, involved in collagen adhesion in the absence of sucrose), *gbpB* (encoding a glucan-binding protein involved in biofilm formation), and *gtfB* (encoding glucosyltransferase B, responsible for sucrose-dependent glucan synthesis and adherence). In *C. albicans*, we quantified the expression of ALS3 (encoding Als3, an adhesin involved in host cell interactions) and the expression of HWP1 (encoding Hwp1, a hyphal wall protein essential for hyphal formation and adhesion). The process involved RNA isolation, cDNA synthesis, and gene expression analysis via qPCR. For this, dual-species biofilms were cultured for 24 hours, washed with PBS, and detached by sonication as previously described. Total RNA was extracted using the Monarch® Total RNA Miniprep Kit (New England Biolabs), following the manufacturer’s instructions. Complementary DNA (cDNA) was synthesized from 10 µL of total RNA using the High-Capacity cDNA Reverse Transcription Kit (Applied Biosystem, US). The reaction mixture included 10 µL of master mix and was incubated in a thermocycler (QuantStudio 3, Thermo Fisher) under the following conditions: 25°C for 10 minutes, 37°C for 120 minutes, 85°C for 5 minutes, and a final hold at 4°C. Negative controls, consisting of RNA samples with all kit reagents except reverse transcriptase, were included to assess potential genomic DNA contamination. The resulting cDNA was stored at −20°C until further use.

Gene-specific primers for *S. mutans* and *C. albicans* were designed based on the *S. mutans* UA159 and *C.albicans* ATCC 90028 genome validated for specificity and efficiency (**Table 1**). Primer specificity was confirmed by melt curve analysis, and efficiency was determined using standard curves with target values of R² ≥ 0.99, efficiency between 90% and 110%, and a slope of approximately −3.3. Each 10 µL qPCR reaction contained 2 µL of cDNA, 8 µL of 2x SYBR Green Supermix (Thermo Fisher), with 1.4µL of each primer, and molecular-grade water. Reactions were performed in triplicate using a QuantStudio 3 thermocycler with the following cycling conditions: initial denaturation at 95°C for 2 minutes, followed by 40 cycles of 95°C for 15 seconds, annealing at primer-specific temperatures (**Table 1**) for 20 seconds, and 72°C for 60 seconds (extension). A melt curve analysis was conducted at the end of the run to confirm the absence of primer-dimers and non-specific amplification. Amplification curves were analyzed using the QuantStudio Design and Analysis software. The threshold was set automatically by the software within the exponential phase of the amplification curves. The resulting Cq values were used for downstream analysis of gene expression levels.

**Table 1:**
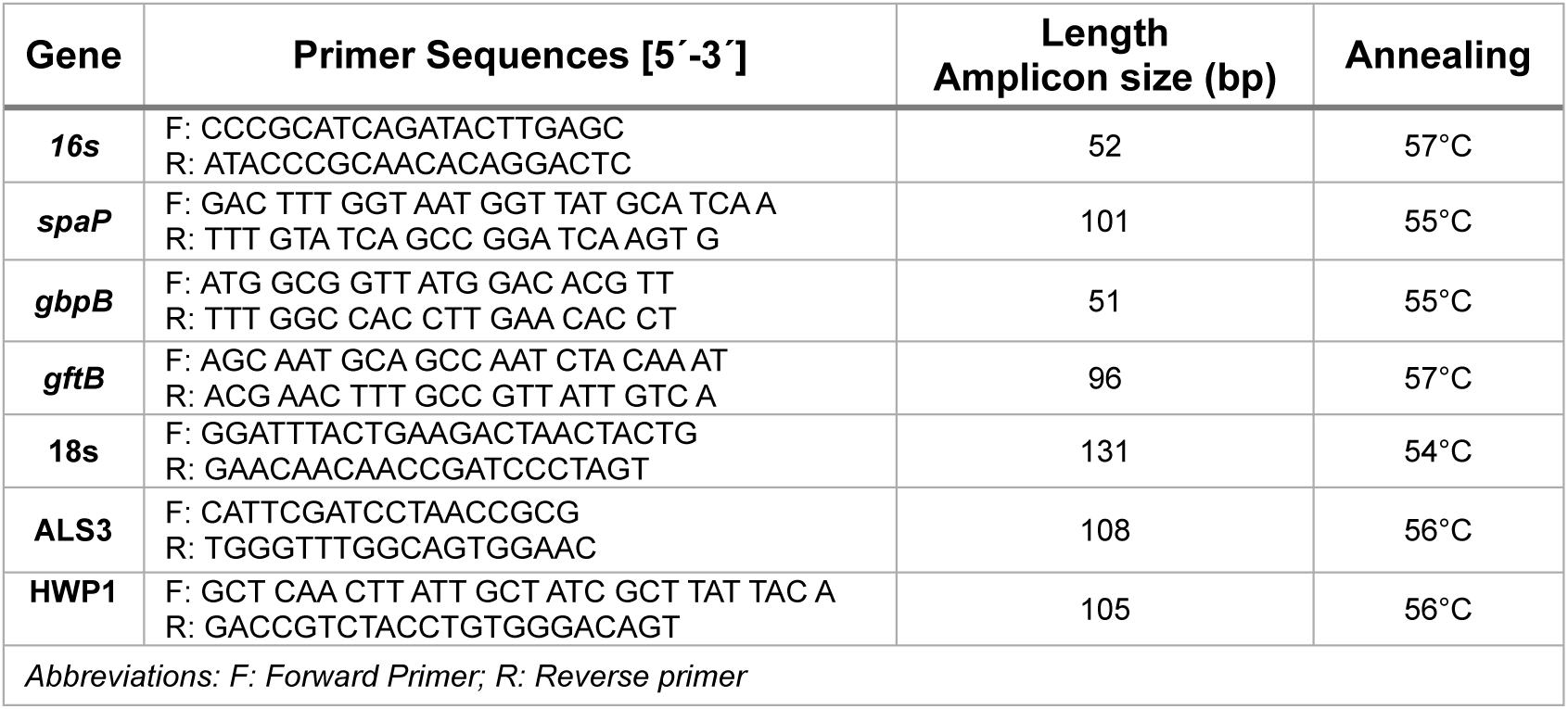
Primer design for Streptococcus mutans and Candida albicans strains.

For *S. mutans* genes, relative expression was normalized to the *16s rRNA* gene, and for *C. albicans* genes, *18s* (a structural ribosome constituent) was selected as the normalizer, supported by their demonstrated stability under similar experimental conditions as reported in the literature (21,30). Relative gene expression was calculated using the ΔΔCt method. Negative controls included no-template controls to rule out contamination.

## 3. Results and discussion

### 3.1. Bioengineered dentin substrates ensure biofilm viability during experimentation

First, we evaluated whether our PDMS-based construct biofilm viability compared to natural dentin. Viability on both substrates was assessed using SYTO9/Propidium Iodide fluorophore labeling at 24 hours, and no significant differences were observed between substrates, and both biofilms exhibited approximately 80% viability (**Figure 2**). The construct consisted of PDMS combined with HAP, collagen, and MGO, which was incorporated to simulate collagen aging through glycation. Although MGO is known to exert some cytotoxic effects on cells (31), its use for surface glycation previous to bacterial inoculation did not impair biofilm viability, probably due to the fact that all unreacted MGO is removed before biofilm experimentation. These results demonstrate the biocompatibility of the bioengineered model to allow biofilm growth in a comparable manner to ex-vivo dentin specimens.

**Figure 2:**
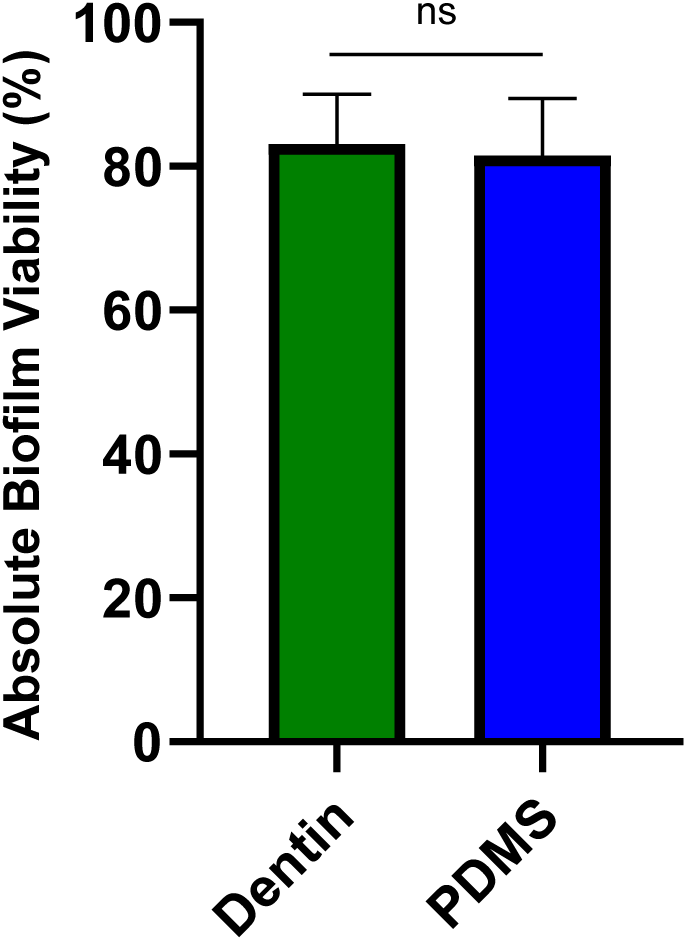
**Cell viability (%) of dual-species *Streptococcus mutans* and *Candida albicans* biofilms on dentin and bioengineered PDMS substrates, assessed using SYTO9 (live cells) and propidium iodide (dead cells) fluorescence after sonication-based detachment**. Biofilms were grown for 24 hours on a 96-well plate. Results are presented as mean values ± standard deviation (SD) from three independent experiments. Statistical analysis was performed using one-way ANOVA with Tukey’s post hoc test. No statistically significant differences were found (p > 0.05).

### 3.2. Biofilm formation on bioengineered constructs closely mimics growth on natural aged dentin

Following CLSM, images of the 24-hour colonized substrates revealed a dense, multilayered, dual-species biofilm structure formed by *S. mutans* and *C. albicans* (**Figure 3**). Adherence to both surfaces was characterized by a dense, compact biofilm, predominantly composed of tightly clustered *S. mutans* cells, primarily concentrated in the layer adjacent to the surface. Furthermore, *S. mutans* displayed the typical microcolony structure associated with local acidification of the biofilm, as expected due to the increase sucrose supplementation in our system(32) *C. albicans* was predominantly observed in hyphal morphology across the z-axis, while EPS was dispersed in small clusters throughout the biofilm. Both substrates exhibited similar features (**Figure 3**) and were comparable to biofilm structures observed on hydroxyapatite substrates, as reported in previous literature (33,34).

**Figure 3:**
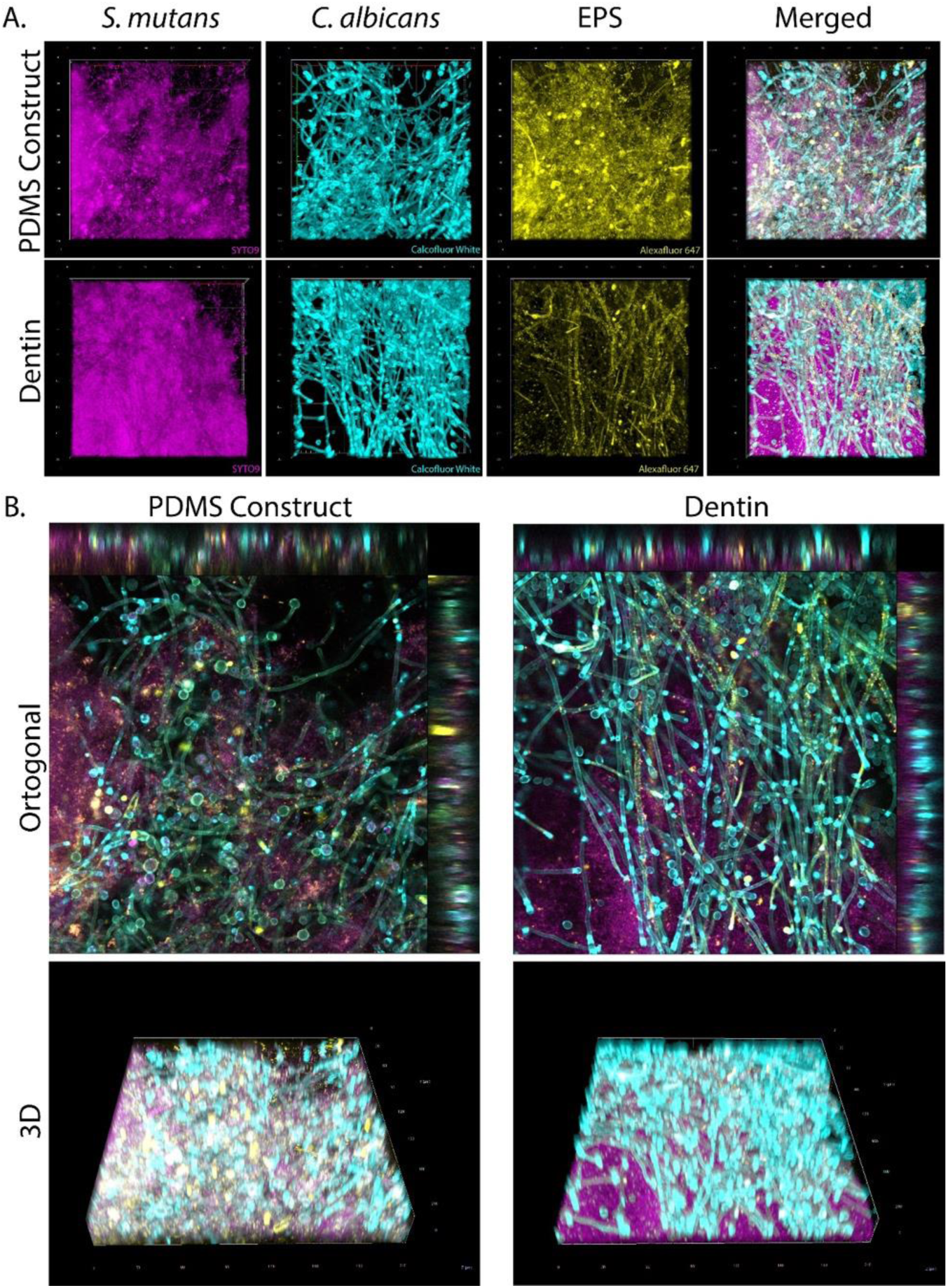
**Confocal laser scanning microscopy (CLSM) images of dual-species biofilms cultured for 24 hours on PDMS and dentin substrates**. *Streptococcus mutans* bacterial microcolonies, stained with SYTO9, are shown in magenta, while fungal cells of *Candida albicans*, stained with Calcofluor White, are displayed in cyan. Extracellular polymeric substances (EPS), labeled with Alexa Fluor 647-dextran, are depicted in yellow. (A) General orthogonal projection (B) 3D up view from individual biofilm components, highlighting the spatial distribution of *S. mutans*, *C. albicans*, and EPS. (C) Composite image combining all three components. (C) Three-dimensional reconstruction of the biofilm, demonstrating its structural complexity (40x magnification).

Furthermore, quantitative analysis of the biofilm images was performed using COMSTAT to evaluate parameters such as biomass, roughness, maximum thickness, average thickness by channel, thickness-to-biomass ratio by channel, and surface-to-biovolume ratio by channel. No significant differences were observed between groups in all the analyzed parameters, confirming the structural similarity of biofilms grown on natural and bioengineered dentin (**Figure 3 and Figure 4**). Maximum biofilm thickness was found not to surpass 40 μm, which is expected given the short incubation times (24 hours) that reflect early biofilm formation(35). Regarding analysis of individual channels, Calcofluor White (*C. albicans)* was observed as the thickest component, whereas SYTO9 (*S. mutans*) represented the thinnest, with both substrates exhibiting similar patterns.

**Figure 4:**
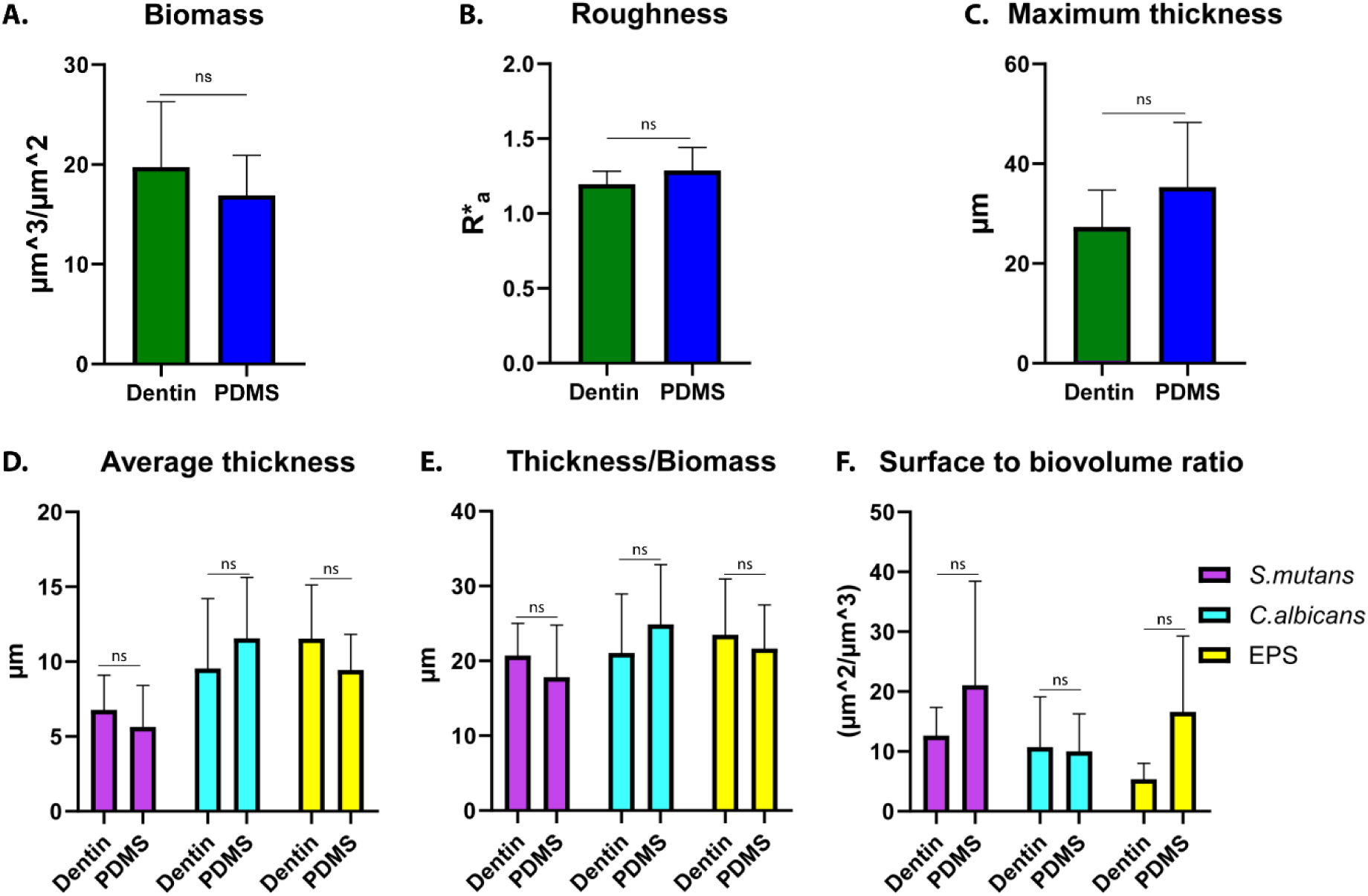
**Quantitative evaluation of biofilm structural and roughness parameters**. Bar graphs illustrate comparisons between ex-vivo dentin and PDMS substrates for total (A) biomass, (B) roughness, and (C) maximum thickness. Biomass was calculated as the sum across channels, and roughness as the average across channels (SYTO9, Calcofluor, Alexa Fluor 647). Additional bar graphs compare dentin and PDMS for each fluorophore label, showing (D) average thickness, (E) thickness-to-biomass ratio, and (F) surface-to-biovolume ratio. Data are presented as mean values ± standard deviations, derived from a minimum of three independent experiments. Statistical analysis was conducted using t-tests and two-way ANOVA, followed by Tukey’s post hoc test. No statistically significant differences were found (p-value >0.05).

A hallmark of oral biofilms is the presence of channels that allow the passage of water and nutrients while dispersing residual metabolites(36,37). In this regard, all components showed a high thickness-to-biomass ratio, suggestive of a porous and dispersed biofilm architecture that is consistent with the presence of channels and areas of lower cellular and EPS density. Surface-to-biovolume analysis revealed greater variability in *S. mutans* and EPS, while *C. albicans* remained consistent across replicates. These findings suggest that *C. albicans* forms a structurally stable and robust framework within the biofilm, whereas *S. mutans* and EPS exhibit higher structural plasticity, likely reflecting adaptive responses to environmental or substrate-related conditions and associated with the formation of microcolony clusters (**Figure 4**). Overall, the present results confirm the structural complexity of dual *S. mutans-C. albicans* biofilms, which is acquired starting the initial stages of biofilm formation (≤ 24 hours) and can further progress at later timepoints (34).

### 3.3. Combined CLSM and fluorometric analysis reveal stable biofilm composition across ex-vivo and bioengineered dentin substrates

To further complement the CLSM observations, we quantified the relative proportion of fluorophores within the biofilm for each experimental condition using a microplate reader (**Figure 5**). To do so, biofilms were detached, resuspended in PBS, and analyzed for fluorophore content. We also assessed whether the absence of salivary pre-coating would alter these proportions. Consistent with CLSM findings, Calcofluor White was the most abundant fluorophore, followed by SYTO9 and, lastly, Alexa Fluor. Interestingly, the EPS signal was found to be lower than the other biofilm components, more closely matching the qualitative observations made from CLSM imaging. Most importantly, similar fluorophore proportions were observed across both substrates, with no significant differences between ex-vivo and bioengineered dentin, confirming the relevance of the *in-vitro* model to accurately represent the *in-vivo* setting. Additionally, the absence of saliva did not significantly impact the relative distribution of fluorophores within the biofilms (**Figure 5**).

**Figure 5:**
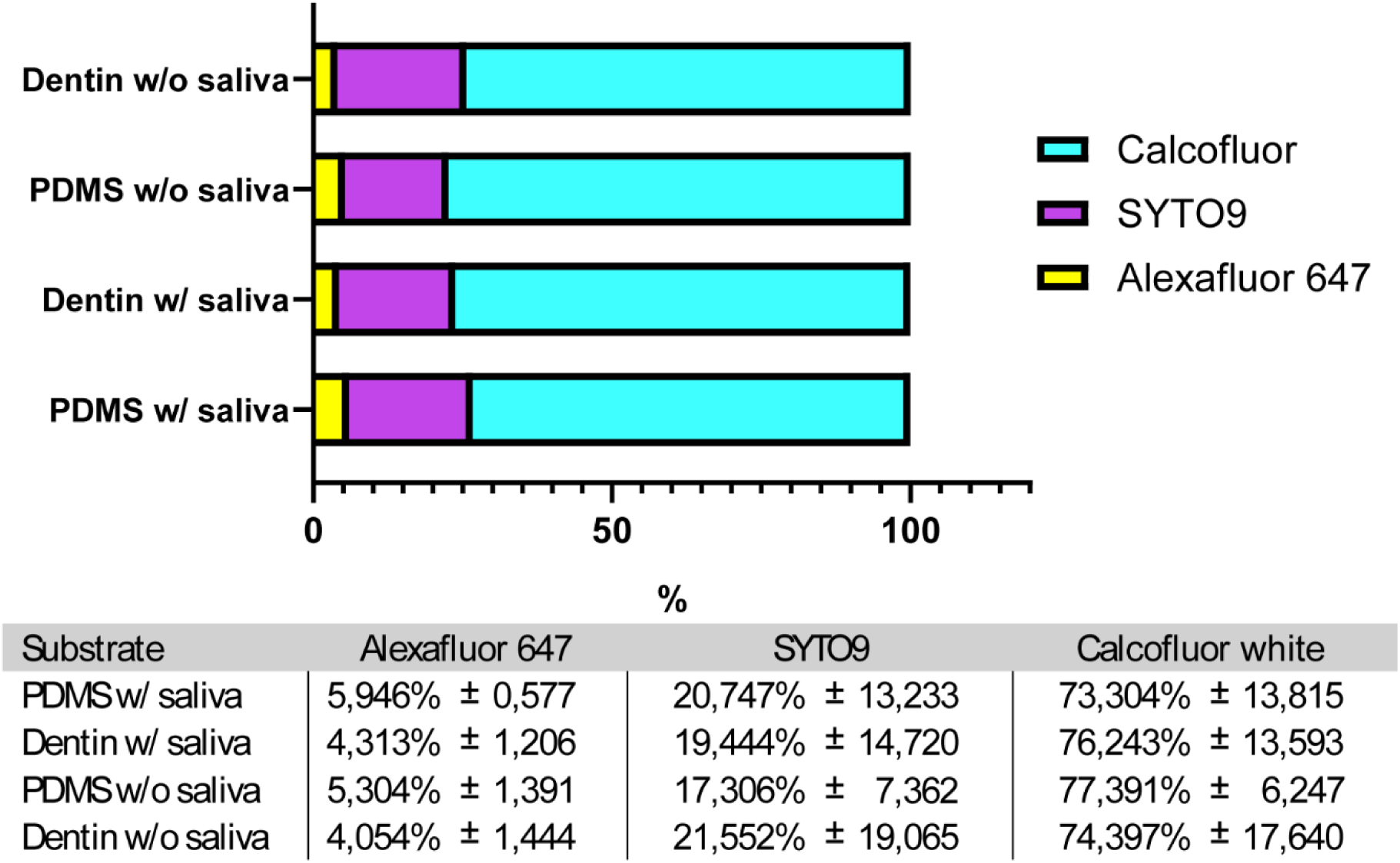
**Proportions of relative fluorescence units (RFU) for SYTO9, Calcofluor White, and Alexa Fluor 647, measured after sonication-based detachment of 24-hour biofilms formed on dentin and PDMS substrate with or without saliva**. Data are expressed as mean values ± standard deviation, calculated from three independent experiments. No statistically significant differences were found with (p > 0.05; two-way ANOVA followed by Tukey’s post hoc test).

### 3.4. Environmental conditions impact cariogenic biofilm virulence on bioengineered dentin-like substrates: a proof of concept

Once the functionality of the bioengineered dentin model to harbor oral biofilms was established, we evaluated if the system allows for the assessment of biofilm virulence changes as a function of environmental conditions in a microscale assay (**Figure 6**). For this, we compared the gene expression of key virulence genes associated with adhesion, biofilm formation, and EPS production in dual-species *S. mutans* and *C. albicans* biofilms grown in the presence and absence of supplemental sucrose.

**Figure 6:**
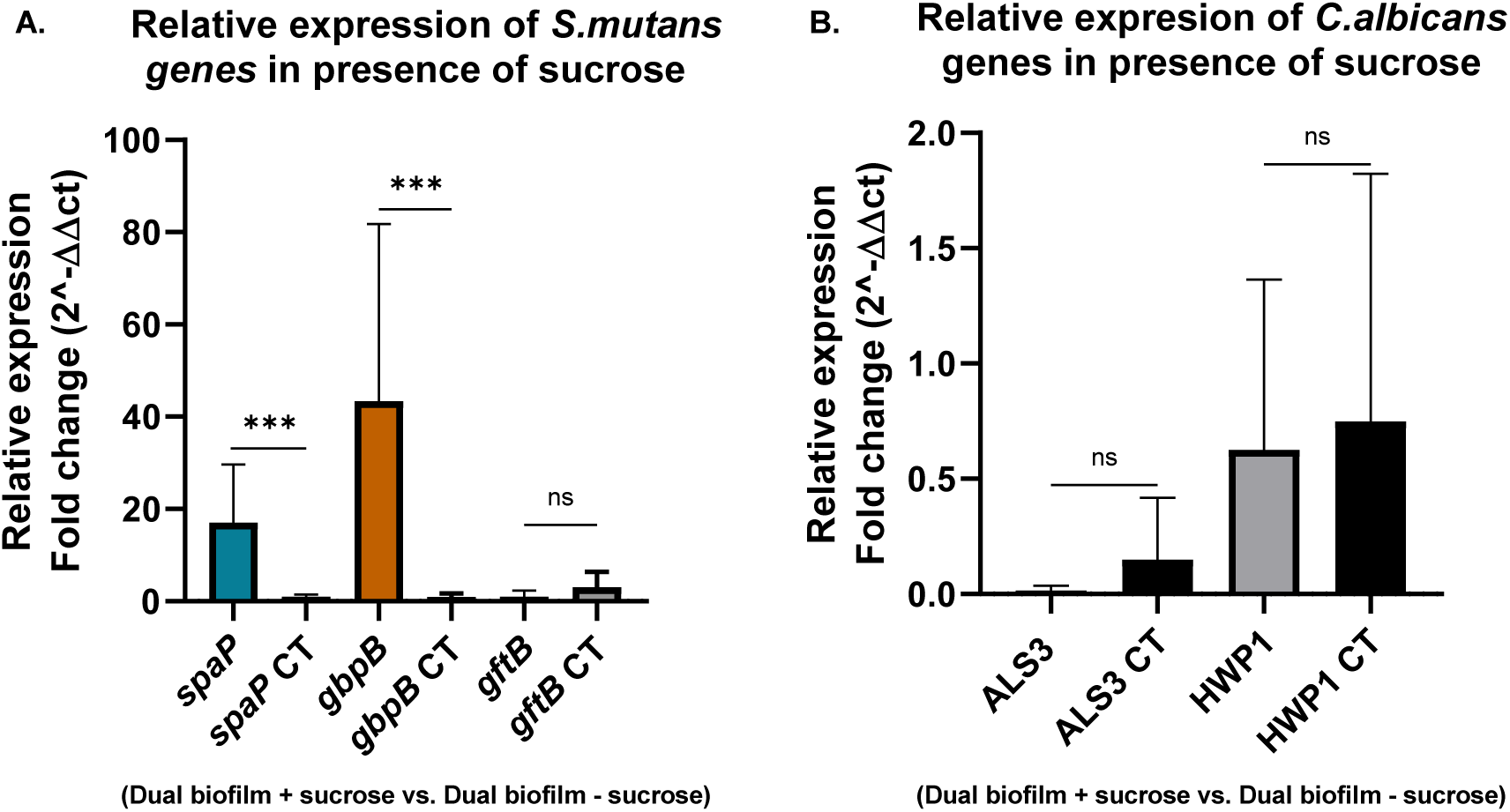
**Expression of *Streptococcus mutans* UA159 genes (*spaP, gbpB, and gtfB*) and *Candida albicans* (*ALS3, HWP1*) analyzed by quantitative real-time PCR**. A) Fold change in *S. mutans* gene expression in a dual-species biofilm in the presence of sucrose. All data were normalized to the 16S housekeeping gene. B) Fold change in *C. albicans* gene expression in the presence of sucrose. Bars labeled “CT” represent the control condition for each gene, used for ΔΔCt calculations. All data were normalized to the 18S housekeeping gene. Results represent at least three independent biological replicates, each performed in triplicate. Statistical analysis was performed using unpaired non-parametric Mann– Whitney U test. Statistically significant differences are indicated with asterisks (*); a p-value less than 0.05 was considered significant.

Our results showed that the sucrose environment significantly upregulates *spaP*, a *S. mutans* gene associated with saliva and collagen binding (38), as well as *gbpB*, which is involved in glucan binding and has been linked to sucrose-enhanced metabolic activity (39). In contrast, *gtfB* was unexpectedly downregulated. We hypothesize that this may be due to the temporal dynamics of gene expression: *gtfB* may exhibits higher transcriptional activity during the exponential stages of biofilm development (4–6 hours), while *gbpB* becomes more dominant at later stages (e.g., 24 hours). This pattern aligns with previous *S. mutans* gene expression studies (34,40). Additionally, reduced sucrose levels in the microenvironment could directly suppress *gtfB* expression, as this gene is highly responsive to sucrose concentration (41). In our experiment, the culture medium was not refreshed, which may have led to a rapid depletion of available sucrose and could partly explain the low EPS levels observed in the CLSM images after 24-hour growth (**Figure 2**).

Secondly, we evaluated *C. albicans* gene expression in the dual-species biofilm model and observed no significant differential expression in the presence versus absence of sucrose. This was expected for the target genes (HWP1, ALS3) as they are not directly sucrose responsive (42). However, we hypothesized that sucrose might indirectly modulate their expression through its known stimulatory effect on *S. mutans* gtfB-mediated glucan synthesis (33). The resulting extracellular glucan matrix was anticipated to serve as a surface signal for *C. albicans*, potentially enhancing hyphal development and upregulating hypha-associated adhesins through the MAPK/Cek1 and cAMP-PKA pathways (43,44). However, contrary to what was expected, both ALS3 and HWP1 were downregulated (**Figure 6B**) despite microscopic evidence of robust hyphal formation (**Figure 5**). In this regard, Martorano-Fernandes et al. (45) reported functional redundancy among adhesins in mixed biofilms, with ALS1 and HWP1 compensating for reduced ALS3 activity. Furthermore, the work of Hwang et al. (33) also positions ALS3 as secondary role in mediating S*. mutans-C. albicans* interactions which could explain low expression of ALS3. However, similar to what was observed for *S. mutans* gtfB expression, it is possible that *C. albicans* ALS3 and HWP1 were downregulated after 24-hour growth in a closed system, and that the overexpression is only observed during exponential growth.

Despite the limitations of this study, our results demonstrate the feasibility of assessing gene expression in caries-relevant oral biofilms after 24-hour growth in the bioengineered microscale array. Nonetheless, the confined microenvironment, defined by the small diameter of the constructs (5 mm) and the use of static culture without medium replenishment, likely resulting in early nutrient depletion, potentially well before the 24-hour endpoint. To capture physiologically relevant gene expression patterns, earlier time points (6–8 hours) are more appropriate, as extended incubation may primarily reflect stress-induced responses or advanced stages of biofilm development(46). These findings also highlight the importance of integrating this construct into a microfluidic system with continuous medium exchange that can better replicate the dynamic *in-vivo* conditions and enhance biological relevance of the system and is part of the future work of this project.

## 4. Conclusion

Overall, the bioengineered PDMS dentin construct was found to support the growth of 24-hour dual-species *S. mutans* and *C. albicans* biofilms, closely resembling the structure and composition of biofilm formation on aged natural dentin. No significant changes in biofilm biomass, bacterial/fungal ratios, and EPS production were observed between the different surfaces, which suggests that this *in-vitro* model is a viable alternative for piloting experiments regarding biofilm formation on aged teeth and reducing the need for human or animal samples. Furthermore, the bioengineered constructs allowed for the exploration of gene expression changes in *S. mutans* and *C. albicans* biofilms as a function of environmental conditions such as the presence/absence of sucrose and polymicrobial interactions. However, further studies incorporating additional variables are necessary to achieve an *in-vitro* model that better replicates the *in-vivo*, clinical scenario. Nevertheless, the development of bioengineered artificial dentin models will offer several advantages for studies regarding dental aging including enhanced reproducibility, cost reduction, and the mitigation of ethical concerns associated with the excessive use of animal and human samples.

## Acknowledgements

This work was financial supported by ANID FONDECYT Regular under Grant #1220804. We also wish to express our deep gratitude to members of the Oral Mechanobiology Laboratory at the Instituto de Ingeniería Biológica y Médica (IIBM) of the Pontificia Universidad Católica de Chile for their outstanding technical assistance and insightful contributions that substantially enriched this research.

